# Draft metagenomes of endolithic cyanobacteria and co-habitants from hyper-arid deserts

**DOI:** 10.1101/2021.02.23.432541

**Authors:** Bayleigh Murray, Micah Dailey, Emine Ertekin, Jocelyne DiRuggiero

**Affiliations:** Johns Hopkins University, Department of Biology, Baltimore, Maryland, USA; Johns Hopkins University, Department of Earth and Planetary Sciences, Baltimore, Maryland, USA

## Abstract

Cyanobacteria are essential to microbial communities inhabiting translucent rocks in hyper-arid deserts. Metagenomic studies revealed unique adaptations of these cyanobacteria but validation of the corresponding metabolic pathways remained challenging without access to isolates. Here we present high-quality metagenome assembled genomes for cyanobacteria, and their heterotrophic companions, isolated from endolithic substrates.

---

In the most arid deserts, where environmental conditions are extreme, microbial communities find refuge inside rocks as a survival strategy (1). The rock habitat protects microorganisms from high UV radiation and drastic temperature fluctuations and promotes water retention within the rock matrix (2). Molecular studies of endolithic communities (within rock) revealed ecosystems spanning all domains of life and multiple trophic levels (3–5). The communities are based on the primary production of cyanobacteria, sometime algae, and are constituted of an assemblage of heterotrophic bacteria and/or archaea, and viruses (6–10). Endolithic communities are highly specific to their lithic substrate with fine scale diversification of the microbial reservoir driven by substrate properties (3, 10).

Cyanobacteria inhabiting endolithic substrates in arid deserts are mostly members of the orders *Chroococcales* (*Chroococcidiopsis* and *Gloeocapsa)*, *Nostocales,* and *Oscillatoriales* (1). Metagenomic studies of endolithic communities revealed unique adaptations of these cyanobacteria and a high number of pathways for secondary metabolites, non-ribosomal peptides, and polyketides, are encoded in their genome (7, 10). However, validation of these pathways remained challenging without access to isolates. Here, we present the metagenome assembled genomes (MAGs) of cyanobacteria isolated from endolithic substrates collected in the Atacama and the Negev Deserts (Table 1). Because these isolates are not purified cultures, their companions - heterotrophic bacteria - have also been sequenced.

**Table 1.** Metagenome and MAGs statistics of endolithic cyanobacteria isolates from the Atacama Desert, Chile, and the Negev Desert, Israel

| Sample Name | Substrate | IMG taxon ID | Metag size Mbp | Bin ID | Taxo_Genus | MAGs Complet<sup>1</sup> % | MAGs Contam<sup>2</sup> % | MAGs size Mbp | MAGs Gene Count | MAGs Scaffold Count |
| --- | --- | --- | --- | --- | --- | --- | --- | --- | --- | --- |
| C-VL-3P3 | calcite | 3300039404 | 11 | 3300039404_1 | *Chroococcidiopsis* | 99.48 | 1.63 | 6.6 | 6630 | 157 |
|  |  |  |  | 3300039404_2 | *Deinococcus* | 97.67 | 0.99 | 4.1 | 4214 | 68 |
| G-Km37-3P1 | Gypsum | 3300039405 | 18.2 | 3300039405_1 | *Methylobacterium* | 100 | 0 | 6.9 | 6942 | 64 |
|  |  |  |  | 3300039405_2 | *Deinococcus* | 97.67 | 0.99 | 4.1 | 4212 | 67 |
| G-Km37-3P3 | gypsum | 3300039416 | 42.9 | 3300039416_1 | *Chroococcidiopsis* | 99.48 | 1.63 | 6.6 | 6618 | 153 |
|  |  |  |  | 3300039416_2 | *Deinococcus* | 97.67 | 0.99 | 4.1 | 4213 | 66 |
| G-MTQ-3P1 | gypsum | 3300038622 | 16.2 | 3300038622_1 | *Chroococcidiopsis* | 99.48 | 1.63 | 6.6 | 6601 | 163 |
|  |  |  |  | 3300038622_2 | *Methylobacterium* | 52.45 | 1.25 | 4.2 | 4745 | 668 |
| G-MTQ-3P2 | gypsum | 3300037877 | 9.9 | 3300037877_1 | *Chroococcidiopsis* | 99.48 | 1.63 | 6.6 | 6608 | 161 |
| H-SG-1P1 | gypsum | 3300039034 | 38.8 | 3300039034_1 | *Chroococcidiopsis* | 99.48 | 1.63 | 6.6 | 6605 | 160 |
| H-SG-2P1 | gypsum | 3300039035 | 43.1 | 3300039035_2 | *Chroococcidiopsis* | 99.48 | 1.63 | 6.6 | 6619 | 155 |
|  |  |  |  | 3300039035_3 | *Deinococcus* | 97.67 | 0.99 | 4.1 | 4230 | 70 |
| I-MTQ-2P3 | ignimbrite | 3300039417 | 20 | 3300039417_1 | *Chroococcidiopsis* | 97.11 | 4.52 | 7.6 | 7825 | 531 |
|  |  |  |  | 3300039417_2 | *Deinococcus* | 98.52 | 0.99 | 4.2 | 4428 | 95 |
|  |  |  |  | 3300039417_3 | *Thermomicrobiales* | 63.91 | 1.89 | 2.4 | 2800 | 503 |
| I-MTQ-3P1 | ignimbrite | 3300039418 | 28.2 | 3300039418_3 | *Deinococcus* | 97.67 | 0.99 | 4.1 | 4241 | 70 |
| I-MTQ-3P3 | ignimbrite | 3300039424 | 30.6 | 3300039424_2 | *Aquamicrobium* | 99.59 | 0.75 | 4.4 | 4417 | 7 |
|  |  |  |  | 3300039424_3 | *Deinococcus* | 97.25 | 0.99 | 4.1 | 4315 | 99 |
|  |  |  |  | 3300039424_4 | *Microcella* | 99.38 | 0.25 | 2.5 | 2464 | 5 |
| I-MTQ-4P3 | ignimbrite | 3300039425 | 10.3 | 3300039425_1 | *Deinococcus* | 97.67 | 0.99 | 4.1 | 4237 | 70 |
| S-NGV-2P1 | sandstone | 3300039401 | 43 | 3300039401_1 | *Chroococcidiopsis* | 99.48 | 1.63 | 6.6 | 6617 | 153 |
|  |  |  |  | 3300039401_2 | *Deinococcus* | 97.67 | 0.99 | 4.1 | 4214 | 67 |
| S-NGV-2P2 | sandstone | 3300039032 | 6.8 | 3300039032_1 | *Chroococcidiopsis* | 99.48 | 1.63 | 6.5 | 6596 | 163 |
| S-NGV-3P2 | sandstone | 3300039033 | 6.8 | 3300039033_1 | MQ *Chroococcidiopsis* | 99.48 | 1.63 | 6.6 | 6618 |  |

<sup>1</sup>MAGs (metagenome assembled genome) completion
<sup>2</sup> MAGs contamination

Cyanobacteria isolates were obtained by incubating ground colonized rock samples collected in the Atacama and the Negev Deserts (3, 4) in Bold’s Basal Medium (11) and in BG11 liquid medium (12) for 5 weeks at 25°C, under 24 μmoles photons/m2/s of white light (WL) using Philips Daylight Deluxe Linear Fluorescent T12 40-W Light Bulbs and a combination of neutral density filters (299 1.2ND and 298 0.15ND, Lee Filters, Burbank, CA). Single colonies were then isolated from 1% agar BG11 medium. These were further grown in BG11 liquid medium under WL and total DNA was extracted from cell pellets using the MoBio PowerSoil DNA extraction kit (MoBio Laboratories Inc., Solana Beach, CA). Nextera with Ranger size selected libraries were made with total DNA and sequenced to 2Gb depth using 2×150 nt reads on Illumina NovaSeq at the DOE’s Joint Genome Institute (JGI). Sequence quality control and assembly where carried out with SPAdes (13) and the annotation was perform with the IMG Annotation Pipeline v.5.0.15 at JGI. MetaBAT version 2.12.1 (14), CheckM v1.0.12 (15), and the GTDB database release 86, GTDB-tk version v0.2.2 were used for binning of MAGs.

High quality MAGs of cyanobacteria were recovered from most samples (Table 1) together with heterotrophic bacteria. All cyanobacteria belong to the *Chroococcidiopsis* genus; *Deinococcus* were the most common “companion” but we also found members of the *Proteobacteria*, *Actinobacteria*, and *Chloroflexi*, illustrating the diversity of these communities.

## Data availability

Raw sequencing data are available from the National Centre for Biotechnology Information under Bioproject Accession PRJNA654119-PRJNA654124, PRJNA677471-PRJNA677478. The metagenome co-assembly and functional annotation are available from the JGI Genome Portal under IMG taxon ID reported in Table 1. To obtain cultures of cyanobacteria isolates, please contact Dr. DiRuggiero at.

## Acknowledgements

These sequence data were produced by the US Department of Energy Joint Genome Institute http://www.jgi.doe.gov/ in collaboration with the user community. We thank the following individuals for their support for library preparation, sequencing, and analysis: Marcel Huntemann, Alicia Clum, Brian Foster, Bryce Foster, Simon Roux, Krishnaveni Palaniappan, Neha Varghese, Supratim Mukherjee, T.B.K. Reddy, Chris Daum, Alex Copeland, , I-Min A. Chen, Natalia N. Ivanova, Nikos C. Kyrpides, Miranda Harmon-smith, Emiley A. Eloe-Fadrosh. This work was supported by NSF grant DEB1556574 and NASA grant NNX15AP18G.

